# Spatiotemporal dynamics of cyanobacterium *Dolichospermum lemmermannii* populations in a bloom-prone region of Lake Superior

**DOI:** 10.1101/2024.02.28.582538

**Authors:** Andrew W. Wood, Jake D. Callaghan, Reane Loiselle, Robert M. McManus, Sandra S. Brovold, Holly A. Wellard Kelly, Elizabeth E. Alexson, Robert W. Sterner, Matthew J. Hudson, Cody S. Sheik

**Author notes:** Corresponding author Cody Sheik, Large Lakes Observatory, University of Minnesota Duluth, 2205 E 4th St, Duluth, MN, 55812.

## Abstract

Cyanobacterial Harmful Algal Blooms (cHABs) are increasingly common in marine and freshwater environments, including the Laurentian Great Lakes (LGL). Lake Superior has seen two large-scale cyanobacterial blooms (2012 and 2018) along the Wisconsin shoreline west of the Apostle Islands, caused by the cyanobacterium *Dolichospermum lemmermannii.* The drivers of bloom formation in Lake Superior are not yet certain, with many factors known to trigger blooms elsewhere in the LGL being absent in Lake Superior. Furthermore, little is known about *D. lemmermannii*’s spatial structure or phenology. Thus, we sought to track the seasonal population dynamics of *D. lemmermannii* to shed light on its growth, physiology, and abundance. In 2021, we used 16S rRNA amplicon and shotgun metagenomic sequencing to characterize spatiotemporal patterns of *D. lemmermannii* abundance and diversity along the bloom-prone Wisconsin shore of Lake Superior. In 2022, we performed net tows and direct colony counts in another localized area. No large-scale bloom event was observed during either year, though several smaller blooms were observed. *D. lemmermannii* abundances were low at nearly all sites and sampling times. Spikes in abundance occurred in July and September, particularly near Siskiwit Bay, a hotspot of bloom formation. We also observed a seasonal shift in heterocyte and akinete abundance indicative of late-season nutrient limitation. Most striking was the seasonal turnover of *D. lemmermannii* strains, suggesting strain adaptation to specific environmental conditions. These results offer valuable and actionable insights for managers and provide a foundation for additional work to clarify drivers of bloom formation in Lake Superior.

## Introduction

Microorganisms are integral to all aquatic food webs, as their metabolic activity drives nutrient availability (e.g., nitrogen, phosphorus, or sulfur) and creates labile carbon sources that fuel higher trophic levels, especially in Lake Superior (Biddanda et al., 2001; Munawar et al., 2009). Aquatic microbial communities are often dominated by a few abundant members, with most microbial diversity residing in the rare biosphere, which acts as a vast microbial seed bank (Caporaso et al., 2012). This contributes to long-term community stability and facilitates recovery following disturbance (Shade et al., 2012). Typically, microbial communities are functionally redundant, such that several species or subspecies perform the same metabolic function, i.e., microbial guild (Louca et al., 2018). Within guilds, specialization on specific substrates or environmental conditions can occur, lessening resource competition between species (Qin et al., 2024). Such specialization can also occur within a microbial species as a subspecies or ‘strain.’ Differences between strains often have observed phenotypic or behavioral changes, such as differences in cyanotoxin production (Cai et al., 2023; Dick et al., 2021) or temperature optima (Eren et al., 2013). These changes are often resolvable at the genome level using molecular approaches (Berry et al., 2017; Dick et al., 2021; Eren et al., 2013) and suggest that in the natural environment, population-level variation is prevalent among most species, including bloom-forming cyanobacteria in Lake Superior (Sheik et al., 2022). It is understood that freshwater microbial populations can comprise many strains, each with genomic variation that may be ecologically significant (Berry et al., 2017; Jaspers and Overmann, 2004). For example, strains of a *Synechococcus* sp. are differentially abundant depending on the water column temperature and light conditions (Berry et al., 2017; Dick et al., 2021; Eren et al., 2013). Furthermore, a strain can arise through horizontal gene transfer by acquiring new genes that facilitate growth in non-typical environments (Tanabe et al., 2018). Time-series surveys of microbial populations in marine and freshwater systems show that species or strain abundance is often linked to the phenology of the system, e.g., the response of organisms to changes in season, where a combination of environmental factors are all co-occurring (Caporaso et al., 2012; Eren et al., 2013; Rohwer et al., 2023). This punctuated periodicity in microbial abundance/activity highlights that ecosystem metabolism will be affected by the timing of when microbial guilds or microbial strains’ metabolisms are active (Van de Waal et al., 2011).

Lake Superior is a cold, oligotrophic lake that has seen an unexpected increase in cyanobacterial blooms in recent years (Sterner et al., 2020), especially from Bark Bay to the Apostle Islands (Figure 1). Awareness of such blooms in non-eutrophic and cold ecosystems is growing (Reinl et al., 2023, 2022) and can cause extensive damage to recreation, fisheries, and water quality (Fristachi et al., 2008). Their appearance in Lake Superior presages changes to the ecosystem services Lake Superior provides and has motivated an urgent scientific effort to explain their emergence. For example, a national travel guide recently expressed concern about Lake Superior as a travel destination because of the newly arising blooms (“Fodor’s No List 2024,” 2023). Lake Superior blooms are primarily attributed to *Dolichospermum lemmermannii* (Sterner et al., 2020), a heterocyte-producing cyanobacterium that initial molecular surveys identified at very low abundances at many points throughout Lake Superior (Sheik et al., 2022). Although many bloom-forming cyanobacterial taxa elsewhere in the Laurentian Great Lakes (LGL) are capable of producing highly dangerous cyanotoxins, e.g., microcystin, anatoxin, or saxitoxin (Miller et al., 2017), only anabaenopeptin was detected in the Lake Superior 2018 bloom (Sterner et al., 2020). Additionally, the genes necessary for microcystin production were absent from a near-complete *Dolichospermum* genome recovered from a 2018 bloom (Sheik et al., 2022). However, since the 2018 bloom, microcystins have been reported in several smaller, polycyanobacterial bloom events in harbor and lake waters consisting of *Microcystis, Aphanizomenon,* and *Dolichospermum* species. Nonetheless, the conditions that trigger *Dolichospermum* and these newer blooms remain somewhat enigmatic. The 2018 bloom followed several earlier intense rain events, and the lake post-stratification was warmer than usual (Sterner et al., 2020). Sediment plumes from rainfall events are a common feature of Lake Superior and deliver limiting nutrients like iron and phosphorus (Cooney et al., 2018; Marcarelli et al., 2019; McKinney et al., 2019), and temperature is important for cyanobacterial growth in other systems elsewhere (Huisman et al., 2018). However, for much of the year, Lake Superior lacks many of the conditions most directly linked to bloom formation in the other LGL; it is colder and more oligotrophic than the other lakes (Sterner et al., 2020), and its watershed lacks heavy industrial and agricultural land use that contributes to intense nutrient influxes at river mouths elsewhere (Watson et al., 2016). The blooms in Superior are also qualitatively distinct from elsewhere in the LGL, occurring along exposed shorelines and not restricted to stagnant bays or river outlets (Sterner et al., 2020). These observations suggest that the drivers of bloom formation in Superior differ from other lakes, but efforts to study them are complicated by their infrequency. Thus, disentangling their triggers will require a better understanding of *Dolichospermum’*s ecology.

To identify variables associated with population dynamics and bloom formation, it is important to determine where and when *Dolichospermum* is most abundant. For example, Reinl et al. (2020) hypothesized that cyanobacterial blooms could be initiated by ‘fluvial seeding’, wherein propagules delivered by cold rivers grow rapidly in warmer near-shore waters. However, Sheik et al. (2022) identified several amplicon sequence variants (ASVs), approximate to a strain, within the water column and sediments and detected strain variation within the genome. This suggests that fluvial seeding drives ASV diversity, or ASV diversity in Lake Superior is driven by habitat (i.e., epilimnion vs deep chlorophyll layer) or water temperature, or a combination of the two. This finding highlights the need for a more thorough study that incorporates source tracking to identify the spatial and temporal patterns of genetic diversity of Lake Superior *Dolichospermum.* These efforts may reveal specific strains associated with bloom hotspots or uncover population structure that points to environmental gradients that drive adaptation to the unique environment of Lake Superior. Mapping the genetic landscape of *D. lemmermannii* can illuminate potential propagule sources (e.g., distinct strains clustered at river mouths) and contribute to effective monitoring efforts by identifying strains whose presence is correlated with bloom frequency (Feist and Lance, 2021). Such a map would also provide a framework for future comparative genomic studies to identify the genetic basis of ecological adaptations (e.g., cold-tolerance) that allow previously transient microbes to thrive in Lake Superior, and be useful for projecting future trends as climate change and other anthropogenic impacts intensify.

This study combines 16S rRNA amplicon and shotgun metagenome sequencing to reveal spatiotemporal patterns in *D. lemmermannii* abundance and genetic diversity in Lake Superior. It also includes directly counted colonies and associated variables in a single bloom-prone embayment area. We focus on the south shore - between Duluth, MN and the Apostle Islands, WI - where cyanobacterial blooms have been most common since they were first confirmed in 2012. We sampled for metagenomes during the summer of 2021 and included samples from open water, protected bays, and along transects extending lakeward from river mouths. Interestingly, the western arm of Lake Superior and its watershed were subjected to a drought, with riverine flow into the lake low to non-existent in some places throughout the study, allowing a focused assessment of strains sourced primarily from the lake. Then, in 2022 we performed a complementary study using net tows and direct counting. Precipitation in 2022 was closer to normal. We found peaks of *D. lemmermannii* abundance in late July and late September, with a hotspot in the area surrounding Siskiwit Bay and no apparent pattern of high river-mouth abundance. We use a genome-resolved metagenomic approach to characterize patterns of genetic diversity in *Dolichospermum* over time and space and present evidence of strain turnover over the course of the summer. We then discuss the actionable insights of our results for stakeholders, their implications for algal blooms in Superior, and new questions they raise.

## Methods

### Sample Collection

Sample stations in 2021 generally occurred along transects that began at tributary mouths and extended perpendicular to 4 km offshore. The term “open-shore” refers to sites at 2 km and 4 km from the tributary mouth and not located within an embayment. Thirty-one such stations were sampled along the south shore of Lake Superior, ranging from Park Point, MN, to the mouth of the Bad River, WI (Figure 1, Table S1). Samples from Park Point to Sea Caves were taken aboard the R/V Kingfisher, with samples from Sand River to Bad River collected from a Northland College research vessel approximately every two weeks from late June through late September in the summer of 2021 (skipping early July). River tributary samples west of Sea Caves were collected from surface waters (depth < 0.5 m) using a rope and attached bucket from either a boat, bridge or directly from the shore, with samples east of Sea Caves taken using a carboy directly from the boat. Sample buckets, carboys, and all bottles were pre-rinsed with sample water before collection. All bottles and carboys were then stored in a closed, dark cooler until they could be processed in the lab. Sampling and filtration of water samples for microbial community analysis were handled in two ways. Surface grab samples were either 1) transported in carboys in a cooler to the laboratory, where they were filtered using 47 mm 0.22 μm PES membrane filters and a filter tower setup (western sites), or 2) water was pumped (using a peristaltic pump) through a 0.22 μm PES Sterivex cartridge filter while on the boat then transported back to the lab on ice (eastern sites). Typically, 500 - 1000 mL of surface water was filtered, depending on sample turbidity. All filters were stored at −80 °C until the time of DNA extractions.

**Figure 1.**
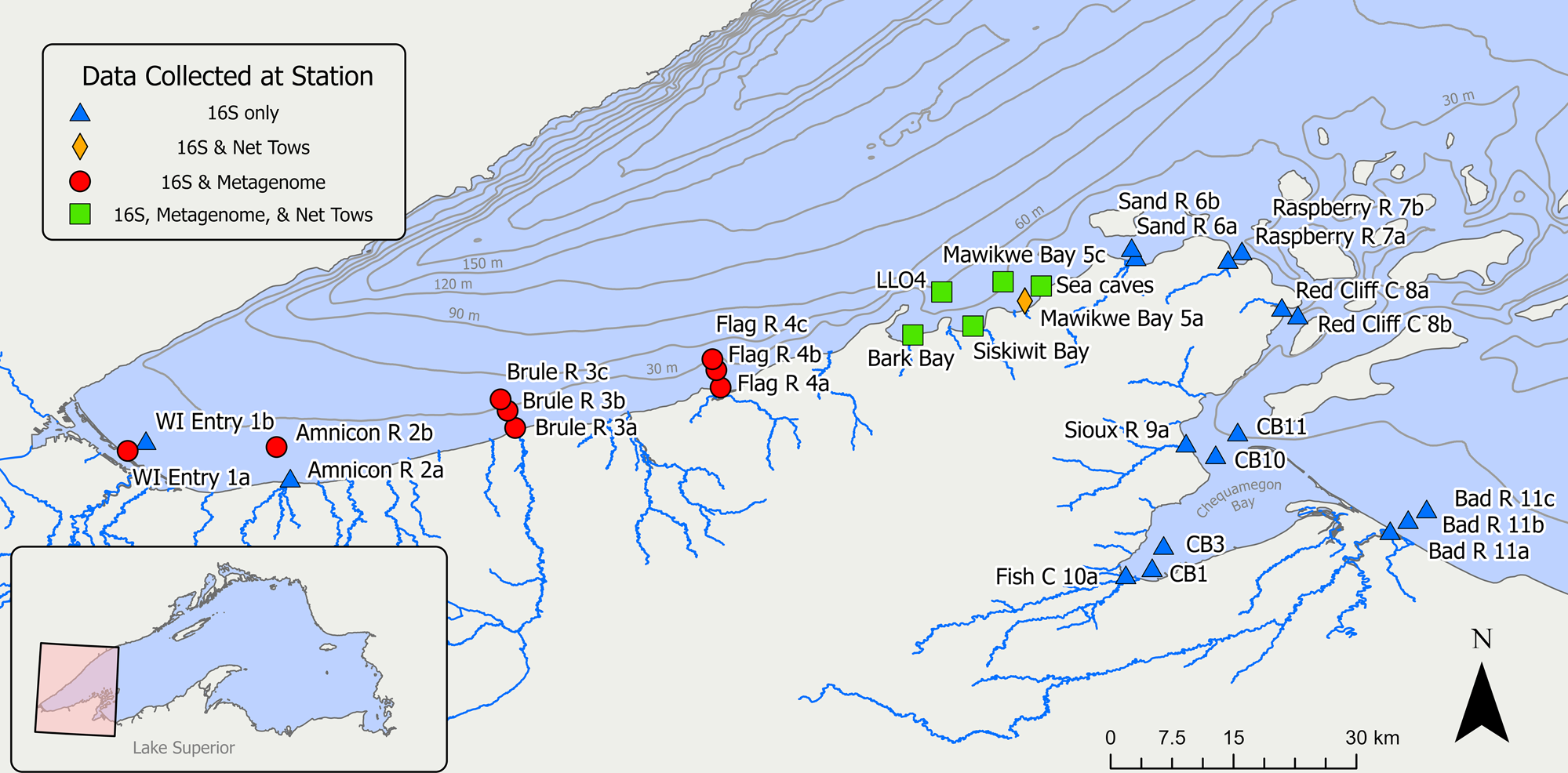
Location of sampling stations from which only 16S rRNA amplicon (blue triangles), both 16S and net tow (gold diamond), both 16S and metagenomic (red circles), or all 16S, metagenomic, and net tow datasets (green squares) were generated. 16S and metagenomic samples were gathered in 2021, while net tows were performed in 2022.

Biweekly vertical zooplankton net tows collections were conducted in 2022 at seven stations (Figure 1, Figure 6). An 80 μm Nitex mesh net with a 50 cm diameter mouth was deployed to a depth of ∼10 m 10x at each site and composited. Samples were then preserved with 1 ml of Lugol’s iodine solution per 100 ml of sample. Enumeration of *D. lemmermannii* colonies began with sample pressurization to collapse cellular aerotopes. 50 ml of sample was then sedimented into 5, 10 ml Üttermohl chambers for over an hour before samples were enumerated microscopically. In addition to counts of the number of colonies, each colony’s total cell count and size were recorded, as well as the presence or absence of heterocytes, akinetes, and *Vorticella spp*. (commonly observed attached to *Dolichospermum* colonies) to better characterize colony state and ecological interactions. Due to uncertainties in the volume of water flowing through each net tow and varying net verticality during windy days, data are presented as relative counts instead of absolute concentrations.

### DNA extraction and sequencing

Sterivex filters were disassembled, and the internal filter housing was aseptically removed. The filter was removed from the cartridge and cut into confetti-sized pieces using a sterile razor blade. DNA was extracted from sample filters and purified with a DNEasy Power Water kit (Qiagen). DNA concentrations were quantified using a Qubit (V3 Invitrogen) and sent to the University of Minnesota Genomics Center for 16S rRNA gene amplicon sequencing. The V4 region of the 16S rRNA gene (515 Forward and 806 Reverse) was sequenced using previously described primer sets (Sheik et al., 2022) with the Illumina MiSeq platform using 2×300 bp paired-end sequencing. Thirty-nine samples were used to create shotgun metagenome libraries using the DNA extracted from the 16S rRNA sequencing. Samples were chosen based on their location and continuity in their time points. Nearshore and offshore samples that successfully amplified for three consecutive time points (e.g., June, July, and August) were kept, resulting in 34 shotgun metagenome libraries. Shotgun metagenome sequencing libraries were created with the Nextera dual-indexed XT library prep kit (Illumina) and were sequenced on a single NovaSeq S4 lane with 2×150 base read lengths.

### Processing of 16S rRNA reads and ASV creation

Reads from 16S rRNA sequence libraries were screened and trimmed of primers using Cutadapt (Martin, 2011). Amplicon Sequence Variants (ASVs) were created using DADA2 (Callahan et al., 2016). ASVs are thought to represent naturally occurring sub-species population variants (Callahan et al., 2019). Briefly, using DADA2, reads were assessed for quality, trimmed, and any PhiX control sequences removed. Forward and reverse reads were de-replicated, ASVs were calculated for forward and reverse sequences, and finally, forward and reverse sequences were merged. Merged ASVs were chimera checked with DADA2. Taxonomic assignment of ASV reads was done using TaxAss v2.1.1 using the custom FreshTrain and Silva databases (Rohwer et al., 2018) and Silva databases (Quast et al., 2013; Yilmaz et al., 2014). ASVs designated as chloroplasts by TaxAss were given further taxonomic assignment using the PR2 database (Guillou et al., 2013).

### Metagenome assembly, genome binning, and taxonomy

Raw reads were first processed using FastP v0.22.0 (Chen et al., 2018), which removes sequencing adapters, trims low-quality regions (Q< 25), and searches and removes any PhiX sequences. Clean reads from each sample were assembled individually or in clusters of five samples to create contigs using MegaHit v1.2.9 (Li et al., 2015). Both assemblies were assessed for the number and quality of Metagenome Assembled Genomes (MAGs) using criteria from Bowers et al. (Bowers et al., 2017). In most cases, the single samples produced better-quality MAGs and were used for downstream analyses. To characterize the metagenomic taxonomic composition, we ran Kraken2 v2.1.3 with default parameters on the final contigs for each sample metagenome (Wood et al., 2019). To create MAGs, we first calculated the relative coverage and abundance of assembled contigs using CoverM *contig* v0.6.1 (Woodcroft, n.d.), mapping reads from all samples to each sample-specific metagenome for maximal resolution. MAGs were generated with Rosella v0.4.2 (Newell, n.d.), Metabat2 (Kang et al., 2019), and SemiBin v1.5.1 (Pan et al., 2022) with default parameters. For MetaBat2 and Rosella, the same Metabat formatted coverage file from CoverM was used. MAGs from the three binning programs were compared using DAStool v1.1.6 (Sieber et al., 2018), with *–search_engine* set to *diamond*, and the unique and highest quality co-occurring MAG was chosen. An additional dereplication step using dRep v3.4.2 (Olm et al., 2017) was used to create a final non-redundant set of MAGs (>95% similarity by ANI). MAG taxonomy was assigned with GTDB-TK v2.3 (Chaumeil et al., 2022), and MAG quality was assessed with CheckM2 v1.0.1 (Chklovski et al., 2023).

### Identification of D. lemmermannii strains

Due to sequencing coverage, no high-quality (>90% completeness and <10% contamination) *Dolichospermum lemmermannii* genomes were recovered. Thus, the high-quality genome from Sheik et al. (2022) was used for downstream analyses. The strain diversity of MAGs was calculated using inStrain v1.7.1 (Olm et al., 2021). InStrain *profile* was used to calculate the within-sample nucleotide diversity of each MAG in our final set, which again contained the Sheik et al. (2022) *Dolichospermum* genome. Each sample’s reads were competitively mapped to this final set of reference MAGs to produce a profile of each MAG for that sample, including average nucleotide identity (ANI) to the reference and major/minor allele frequencies. InStrain *compare* was then used to calculate ANI in all pairwise comparisons of the 34 profiles generated in the previous step and to generate a dendrogram.

## Results

### 16S rRNA gene amplicon sequencing

Using a high throughput 16S rRNA gene sequencing approach, four *D. lemmermannii* Amplicon Sequence Variants (ASVs) were identified in our dataset (Figure 2). These sequences were all highly similar (> 98% similar by BLASTn) to the 16S ribosomal gene identified in the 2018 *D. lemmermannii* bloom (Sheik et al., 2022). The ASVs’ relative abundance was generally low (< 1.5% of the total sequences recovered) and varied in abundance from open-shore to near-shore Lake Superior stations. ASV_405 and ASV_964 were the most abundant and frequently encountered, especially in late July and September. In June, we only detected ASV_405 and ASV_964 but primarily in stations near or within Chequamegon Bay. Due to timing, we were unable to sample during early July. However, in Late July, we saw an increased presence of ASV_405, ASV_677, and ASV_964 in open-shore stations from the Lake Superior entrance to the Sea Caves. ASV_677 was more consistently found in these stations than ASV_405 and ASV_964, which consistently varied between Wisconsin Entry and Flag stations. However, from Bark Bay to the Sea Caves, the abundance of these three ASVs was consistently high. The abundance of these ASVs coincides with several small blooms observed from July 18 near the Sea Caves to July 21st from the Lake Superior shoreline at Park Point, Duluth, MN. Pictures of the cyanobacterium from Park Point are consistent with *D. lemmermmanni*’s bloom presence and cellular phenotype (Figure 3A-D). Interestingly, these ASVs were typically more abundant in stations further from shore, but there was great station-to-station variation (Figure 1, “a” is closest to the tributary outlet, while “c” is furthest; see Table S1). In both August samples, *D. lemmermannii* ASVs were either at very low abundance or not present in our samples. In early September, two ASVs were present, ASV_677 was observed in stations near the Wisconsin entry, while ASV_405 was in Chequamegon Bay. In late September, ASV_405 was observed from Wisconsin Point to the Sea Caves and may constitute a second small bloom. Despite two potential blooms being identified during this time, we found weak positive but nonsignificant correlations between ASV abundance and either water geochemistry or physical properties of the water column, e.g., temperature, depth or time of sampling (Table S2).

**Figure 2.**
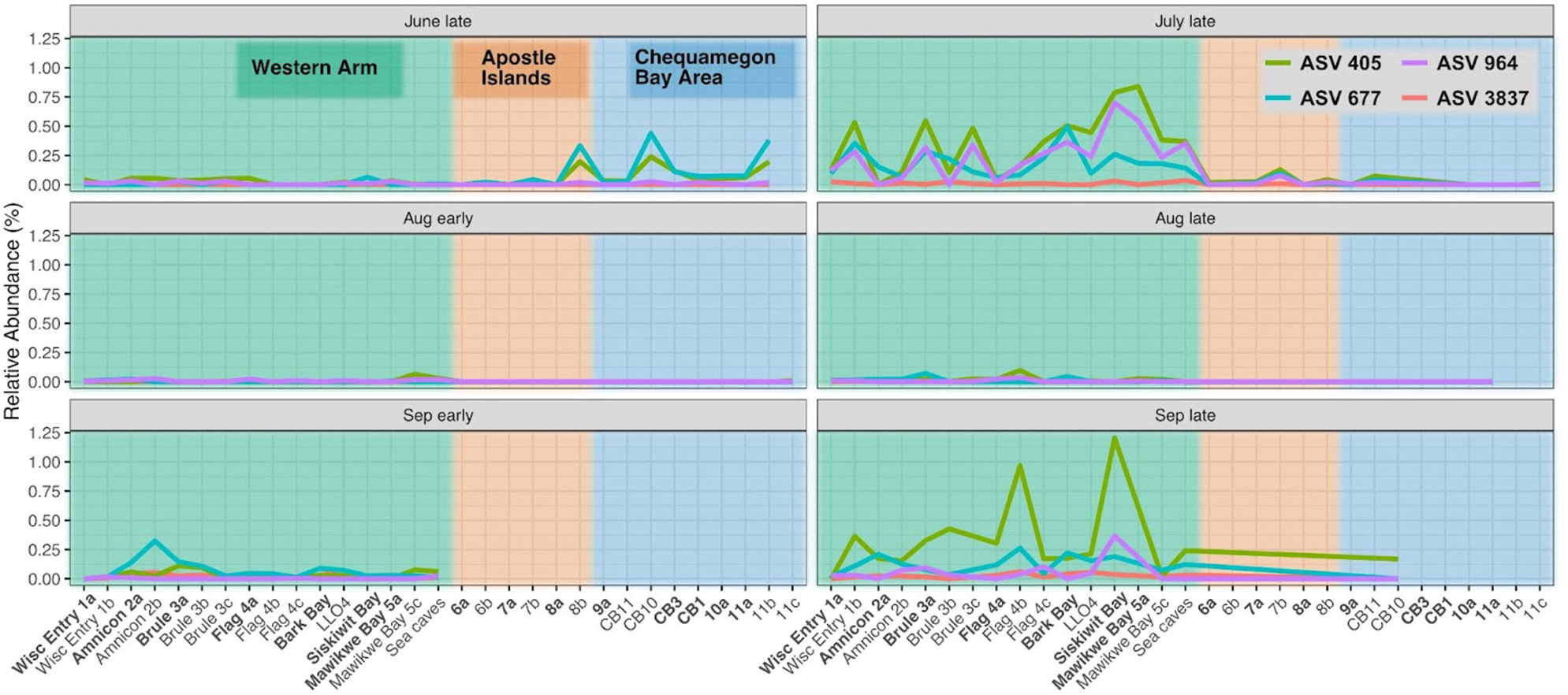
The relative abundance of *D. lemmermannii* Amplicon Sequence Variants (ASVs) identified during the sampling time course. Background colors correspond to the location of sampling stations. From left to right, they are the open shore Western Arm (Green, Wisconsin Entry 1 - Sea Caves), Apostle Islands (Orange, stations 6a - 8b), and the Chequamegon Bay Area (Blue, 9a - 11c). Station names in bold indicate samples taken at tributary mouths.

**Figure 3.**
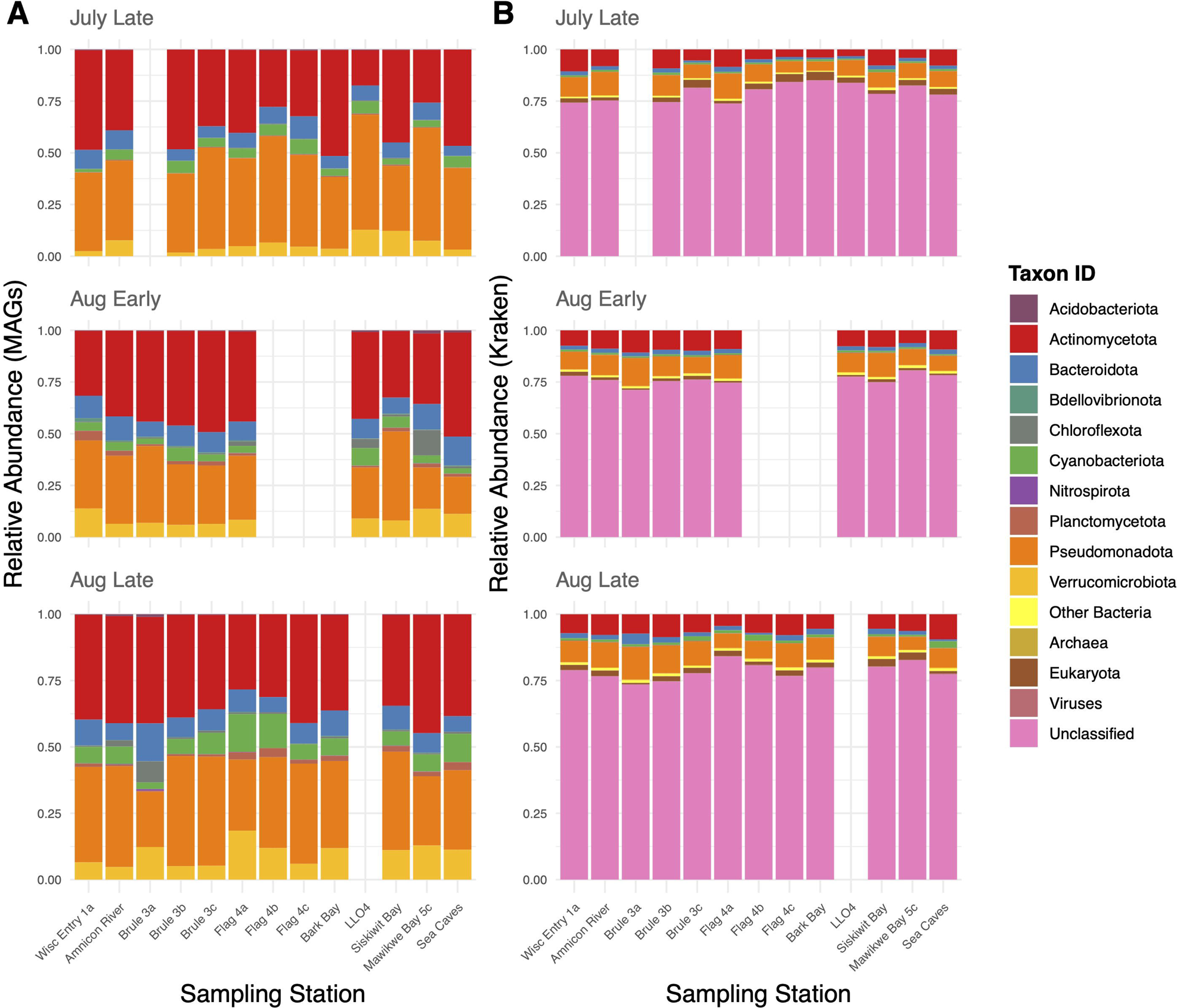
Images of a minor, short-lived bloom in Lake Superior along the Park Point Shoreline on July 21, 2021. A) The bloom from the shore, B) observed cyanobacterial colonies from a sample taken from the bloom, and C and D) images of two colonies observed at 400x.

### Metagenome assembly, genome binning, and abundance

After screening for quality and completeness, 1,364 bins were recovered from the thirty-four samples sequenced for shotgun metagenomics. Dereplication (i.e., clustering highly similar genomes to a single representative genome) reduced the total number of genomes to a final non-redundant set of 112 high-quality MAGs. None of these were identified as *Dolichospermum spp.*. Still, inspection of the raw bins generated from each sample (i.e., before filtering for completeness and quality with DAStool) revealed partial *Dolichospermum* genomes in three samples, all collected on July 20th, 2021: LLO4 (60% complete), Sea Caves (15% complete), and Siskiwit Bay (41% complete). The recovery and presence of partial genomes are likely due to the low abundance of this organism at the time of sampling and the sequencing depth, which is the number of sequences recovered per sample. By sequencing more, we would likely have recovered near-complete genomes. Furthermore, because water samples were not pre-filtered and instead concentrated onto 0.22 μm filters, a large proportion of particle-associated and eukaryotic taxa likely account for many unmapped reads. Indeed, most metagenomic reads for each sample did not map to any reference MAG (Figure 4A; mean unmapped: 65.6% and standard deviation: 6.1%) The taxonomic assignment of the assembled contigs shows similar results and highlights the large number of unidentified contigs, that are likely from Eukaryotes that have yet to be genome-sequenced and incorporated into Kraken2 databases (Figure 4B).

**Figure 4.**
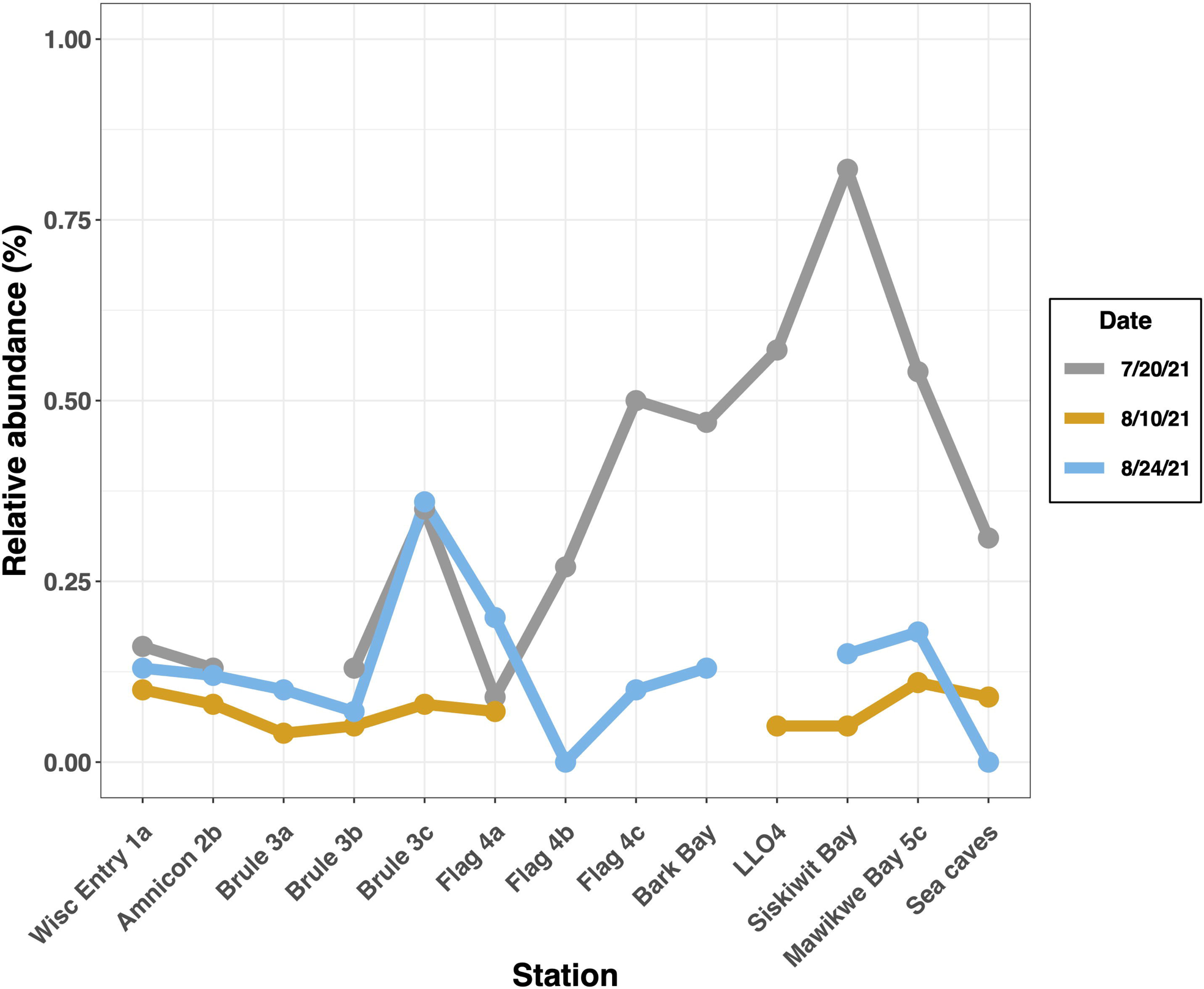
The abundance of sequencing reads mapped to MAGs with taxonomy assigned by GTDB (A) and the taxonomy of assembled contigs using Kraken2 (B). Empty columns represent samples that failed to sequence. To clearly see the abundance of MAGs, reads not mapping to any MAG (mean unmapped: 65.6% and standard deviation: 6.1%) are not shown in (A), and abundances of mapped reads are scaled to 1. The category “Other Bacteria” in (B) represents phyla at extremely low abundance.

Despite not recovering a complete genome from this sequencing effort, we used the nearly complete *D. lemmermannii* genome from Sheik et al. (2022) for inStrain analyses, as our recovered low-quality MAGs were highly similar by average nucleotide identity (ANI > 96). *D. lemmermannii* was detected in all but two samples (Sea Caves and Flag River 4b, both from August 24th) after mapping metagenomic reads to these reference MAGs with CoverM (Figure 5). Like the 16S rRNA amplicons, the average relative abundance of *D. lemmermannii* using the mapped fraction of reads was generally low, 0.19%, with a maximum of 0.82% in Siskiwit Bay on July 20th. *D. lemmermannii* was most abundant on July 20th at nearly all sites, with the highest abundances recorded at sites surrounding Siskiwit Bay, corroborating patterns in the 16S rRNA amplicon data (Figure 2).

**Figure 5.**
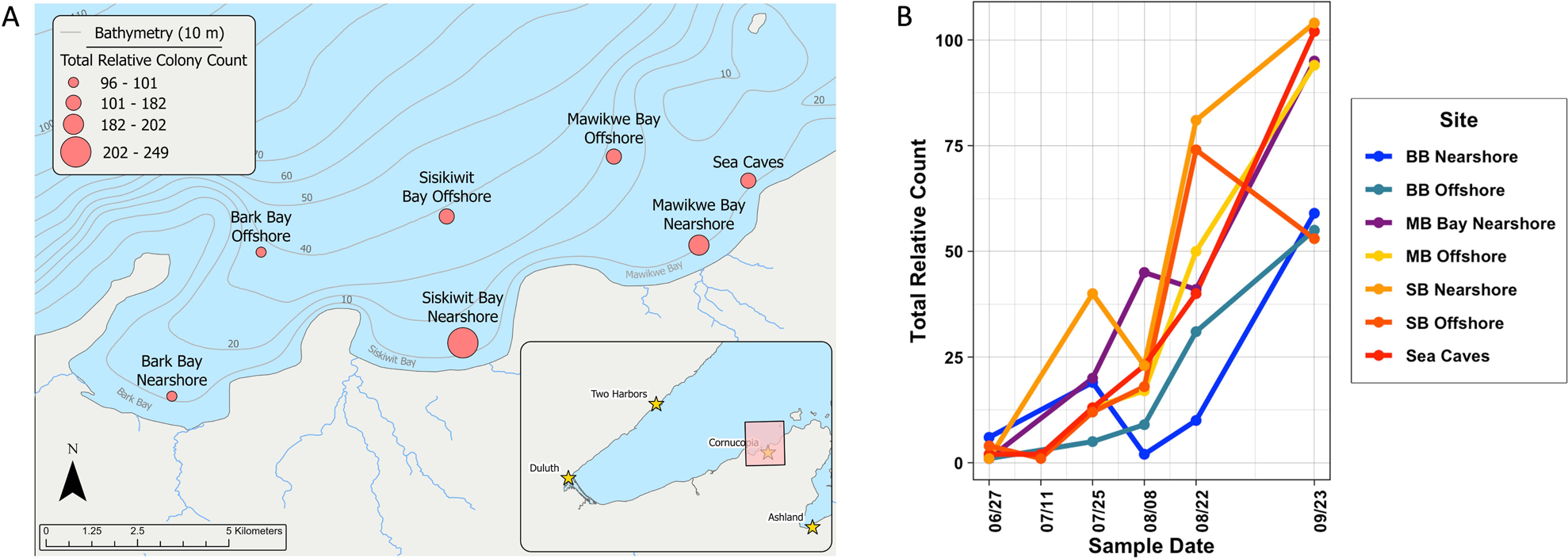
Total relative abundance of *D. lemmermannii* at stations on each sample date. The Relative abundance was calculated with CoverM based on the mapped fraction of reads to final reference MAGs.

### Dolichospermum direct counts

*D. lemmermannii* relative abundances displayed substantial spatial and temporal variability among the seven sites sampled. Spatially, sites less than 1 km from shore exhibited higher colony counts than offshore sites within the same bay. Additionally, Sea Caves, Siskiwit Bay, and Mawikwe Bay consistently had higher total *D. lemmermannii* counts than both Bark Bay sites (Figure 6A). Between June and September, all sites experienced a five to ten-fold increase in total colonies, most of which occurred between the August and September sampling dates (Figure 6B). Colony size did not vary significantly with time (one-way ANOVA on log-transformed data, p = 0.11) (Table 1). Heterocytes were common but variable through time. Akinete presence exhibited seasonality, with higher prevalence later in the year than earlier. Finally, the co-occurrence of *Vorticella spp.* anchored onto *D. lemmermannii* was the rule, with occurrence frequencies ranging between 73-100% of the colonies throughout the sampling period.

**Figure 6.**
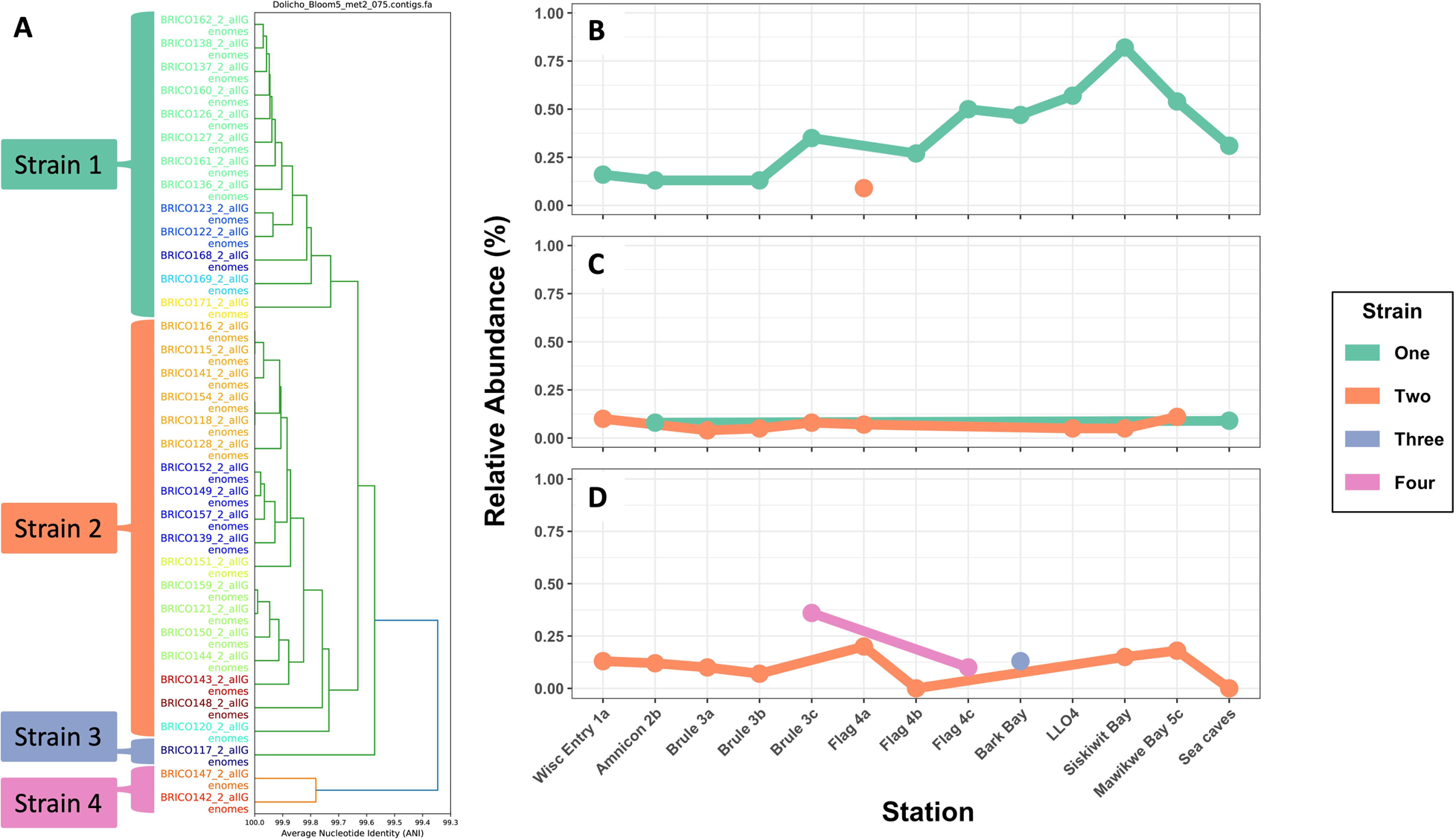
Detailed map of net tow sites with relative *D. lemmermannii* colony counts totaled across sampling dates (A). Circles scaled by count. Total relative colony counts at each date (B) with lines colored by sample site.

**Table 1.**
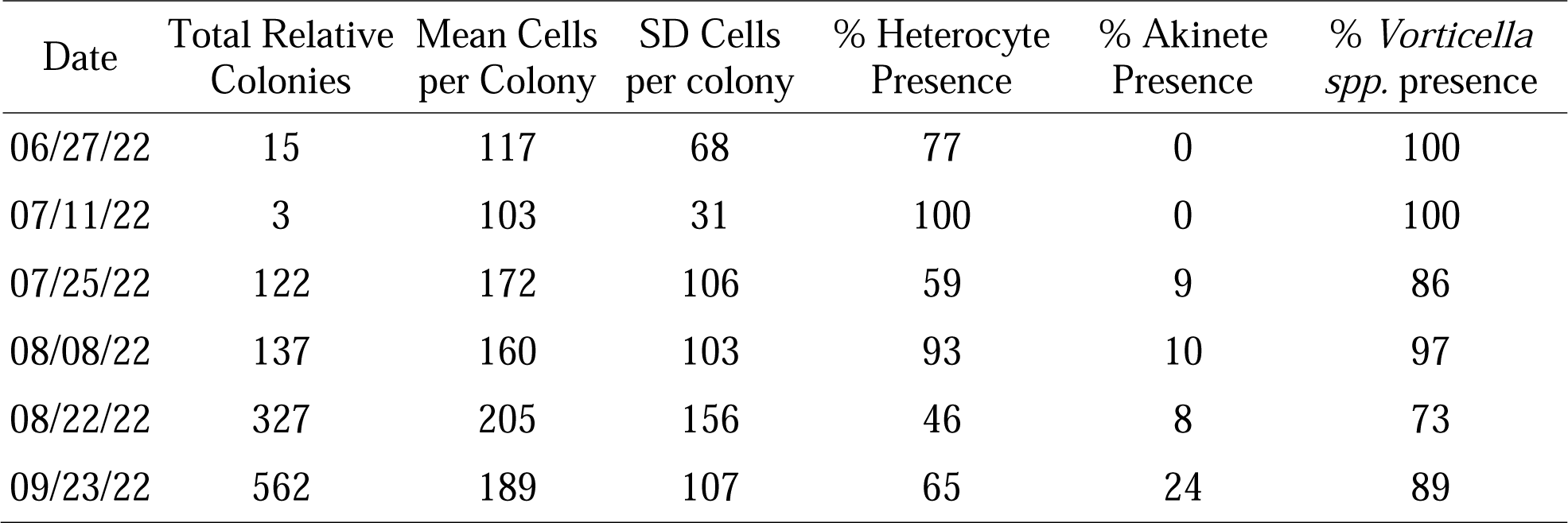
*D. lemmermannii* colony counts from 2022 net tows, including heterocyte, akinete, and *Vorticella spp.* presence.

### Identification of D. lemmermannii strains

The inStrain profiles of the *D. lemmermannii* reference MAG were recovered from all 34 samples and show that sequences related to *D. lemmermannii* were present in each sample and were highly similar (>98% genome-wide Average Nucleotide Identity (ANI)) to the reference genome. For comparison, ANI >95% is considered to be from the same species (Olm et al., 2017). Pairwise comparisons of all profiles to each other revealed similarly high ANI (i.e,. >98%) among all samples. However, no two samples had an ANI of >99.999%, which is the inStrain threshold to designate profiles as having no fixed differences, i.e., the strains are identical (Olm et al., 2021). When clustered by ANI similarity and clusters visualized via dendrogram, four clades are apparent, which we designate as strains: two abundant sampling strains, Strain-1 and -2, and two minor strains, Strain-3 and -4 (Figure 7A). Strain-3 was identified at only one sample time (Strain 3 in Figure 7A). By labeling our sampling map at each time point according to strain present in each sample (Figure 8), we identified a temporal population structure. Strain-1 was the primary strain on the July 20th sampling, with only one sample identified as Strain-2 (Figure 7B, Figure 8A). On August 10th, Strain-1 was mostly displaced by Strain-2 (Figure 7C, Figure 8B), and by August 24th Strain-2 was dominant with Strains-3 & -4 appearing on this final sampling date and Strain-1 entirely absent (Figure 7D, Figure 8C).

**Figure 7.**
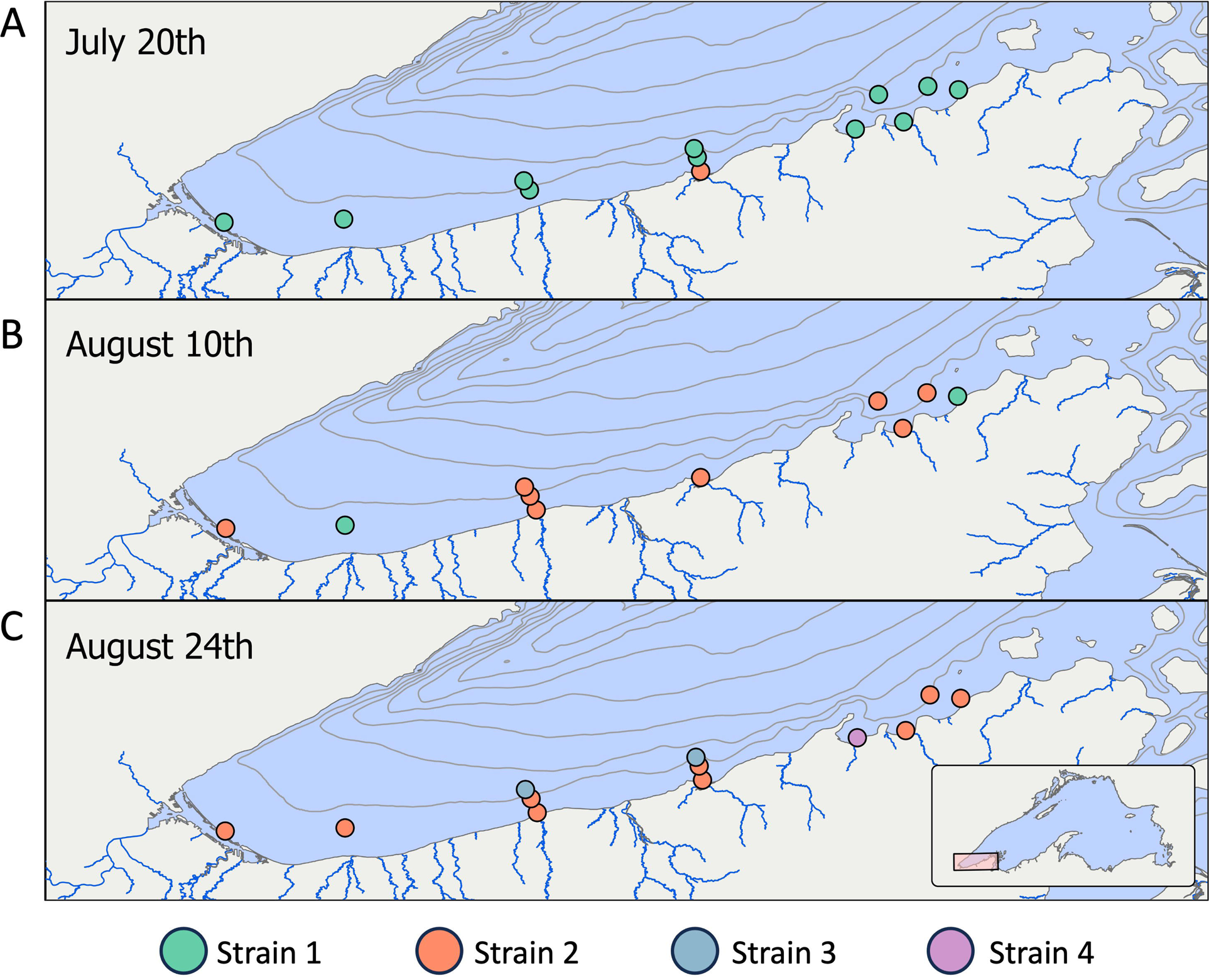
*D. lemmermannii* strain characterization. (A) Dendrogram clustering *D. lemmermannii* by ANI calculated from inStrain MAG profiles of each sample. Colored brackets indicate strain membership. Strain relative abundance at each sample site on (B) July 20th, (C) August 10th, and (D) August 24th.

**Figure 8.**
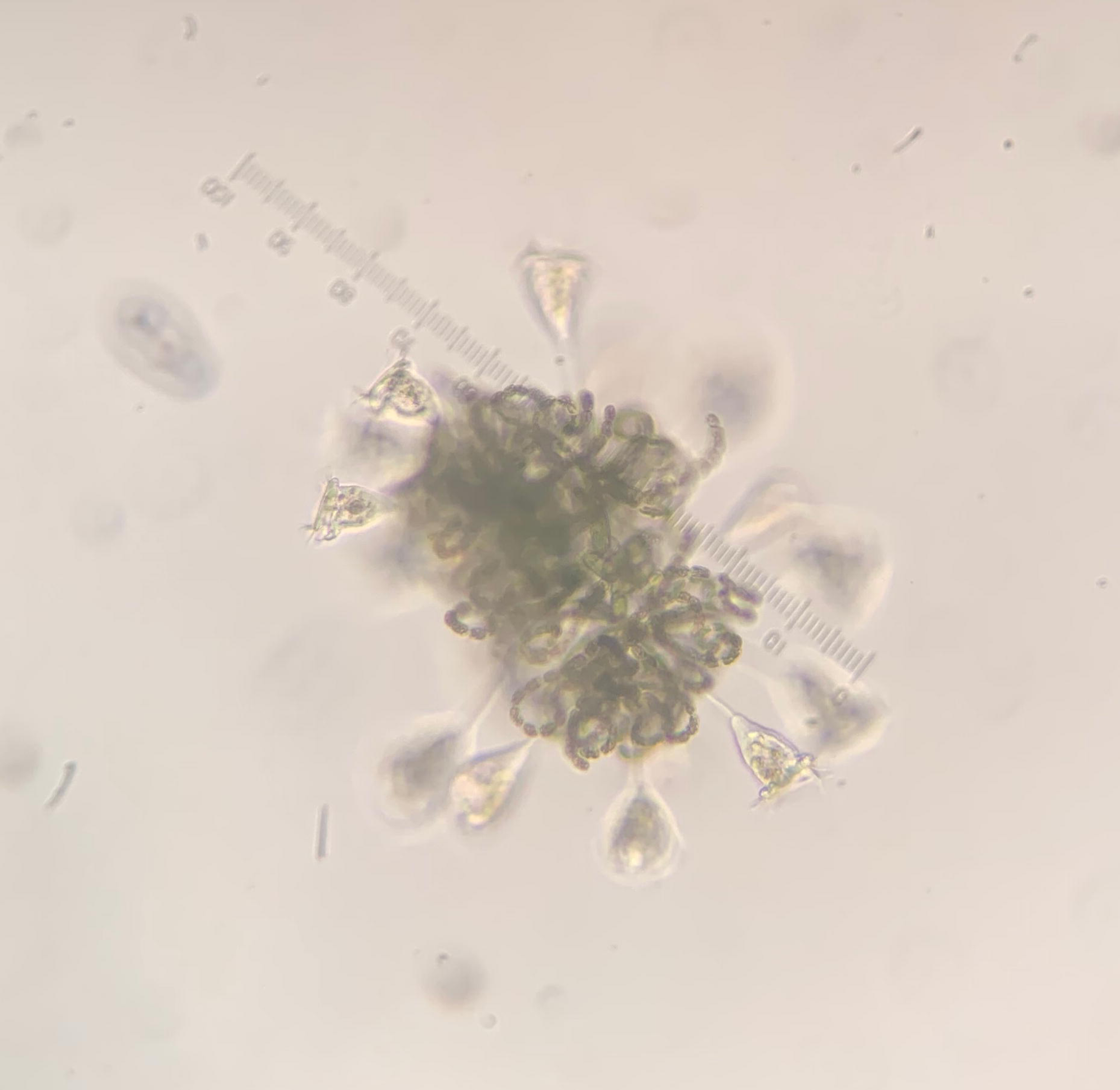
Presence and absence of *D. lemmermannii* strains at each sample site on (A) July 20th, (B) August 10th, and (C) August 24th.

## Discussion

It has been well understood that populations of organisms are genetic amalgams, and when compared at the individual level, variation between genomes can be quantified (Lenski, 2017). These subtle variations among individuals can manifest phenotypic differences that allow subpopulations to specialize within their ecological niche (Eren et al., 2013). As the quality and quantity of DNA sequences have improved, so too has the ability to evaluate genotypes at the population level (Olm et al., 2021; Truong et al., 2017; Zhao et al., 2023). These methods can also facilitate environmental source tracking (Sheik et al., 2018), where it is possible to identify where individual populations are arriving from (Han et al., 2020; Knights et al., 2011). In Lake Superior, recurring cyanobacterial blooms are counterintuitive, as extreme oligotrophic conditions and cold temperatures are thought to discourage bloom formation (Reinl et al., 2023, 2022; Sterner et al., 2020). Previous work has suggested that riverine inputs may be the source of these cyanobacteria (Reinl et al., 2020). However, 16S rRNA gene amplicons suggest several populations exist in the epilimnion of Lake Superior (Sheik et al., 2022) that could be sourced from allochthonous or autochthonous populations, further complicating efforts to identify drivers of these blooms.

Studying the ecology of *D. lemmermannii* in Lake Superior has been challenging, as this organism does not bloom consistently and has resisted cultivation. Population abundances in the lake are also often low, and previous work has relied on sparse data from individual blooms or a single season (Sheik et al., 2022; Sterner et al., 2020). To now, our knowledge of the spatial and temporal variation of *D. lemmermannii* has been very limited. The present study’s detailed sampling and genomic methodology provide an unprecedented view of *D. lemmermannii*’s phenology on a long stretch of bloom-prone shoreline. We found that *D. lemmermannii* is nearly ubiquitous even in non-bloom water samples along the south shore of Lake Superior, but with higher relative abundance (observed in both the metagenome and 16S rRNA amplicon datasets) during mid-to-late summer. A seasonal buildup of *D. lemmermannii* was further corroborated by the 2022 net tows. These dynamics are also consistent with recent findings that *D. lemmermannii* is present in Lake Superior’s epilimnion and deep chlorophyll maximum in late July but not early in the season (Sheik et al., 2022). Our results, however, do not fully reject the fluvial seeding hypothesis of bloom formation (Reinl et al., 2020), as abundance patterns from nearshore and offshore sampling transects show variable abundances of *D. lemmermannii* ASVs. In several instances, we found that *D. lemmermannii* was abundant offshore and near the river outflows (See Figure 2, comparing samples near shore, labeled with an A, to offshore samples, labeled with a B and C). That said, our sampling occurred during a generally low-precipitation year and did not coincide with large precipitation events when nutrient influxes and river outflows would be most intense. Thus, our data likely represent a baseline expectation for population dynamics, and a multi-year sampling and sequencing effort targeting inland and offshore *D. lemmermannii* populations would be necessary to test this hypothesis more directly.

The temporal structure observed in our genomic diversity data from July through August (Figure 7B-D, Figure 8) points to a seasonal succession of *D. lemmermannii* strains. This may be an important finding for clarifying our understanding of bloom dynamics because even highly similar cyanobacterial genomes can be adapted to different ecological conditions and occupy distinct niches, with implications for community composition and function, and even for toxin-producing potential (Österholm et al., 2020; Van de Waal et al., 2011). In Florida Bay, for example, harmful algal bloom formation is associated with strain succession of the bloom-forming cyanobacterium *Synechoccous elongatus*, demonstrating how impactful small genomic differences can be to ecosystem ecology (Berry et al., 2015). Bloom formation depends on the intersection of the right environmental conditions with the bloom-forming organism. If *D. lemmermannii* strains are adapted to distinct ecological niches and dominate the community at different times, mismatches between ideal conditions and strain ecology may determine if blooms form.

What then drives strain succession, and where do different strains originate in Lake Superior? As with the abundance data, the pattern of strain succession doesn’t directly implicate fluvial seeding (Reinl et al., 2020) as a source of novel strains, but neither does it reject this hypothesis, and the same caveats regarding low rainfall in 2021 apply here. Profiling the microbial communities of the rivers and their source lakes would be an additional improvement that would allow us to test this hypothesis. Another possibility is that in non-blooming years, they reside in the rare but active portion of the microbial biosphere and emerge as environmental conditions select for strains with different adaptations (Caporaso et al., 2012). The diverse 16S ASVs detected by Sheik et al. (2022) throughout Lake Superior in the water column and sediment are consistent with this hypothesis. However, with no correlation detected between the environmental variables we measured and *D. lemmermannii* dynamics, it is unclear what conditions might be driving this selection. Future research should identify these variables and determine if ideal bloom conditions exist for long or short time windows.

Other seasonal patterns of potential relevance to blooms pertain to cell differentiation and with close associations with other organisms. Heterocytes (e.g., specialized cells where nitrogen fixation can occur) are typically produced under N-poor conditions, and the role of akinetes in year-to-year carryover of populations is still unknown. Studies of cross-domain microbial consortia are still in the early stages, but identifying closely interacting species, like cyanobacteria and protozoa, suggests further studies are necessary. Our data provide a first inkling into the seasonal dynamics of these specialized cells and the common presence of a protozoan closely associated with *D. lemmermannii* colonies. Heterocytes were variable from sampling to sampling with no obvious strong seasonal pattern. Lake Superior has year-round excess nitrate, but whether individual species rely more heavily on reduced forms of N, like ammonium, is still poorly understood. Environmental factors can trigger akinete differentiation within the *Nostocaceae* family, including low temperature, low light, and various nutrient limitations such as phosphate, K+ ions, or iron (Hori et al., 2003; Kaplan-Levy et al., 2010; Meeks and Elhai, 2002). Although we cannot determine what stimulated akinete differentiation from our data, we detected significant, proportional increases during late September (Table 1). This shift in heterocyte and akinete abundance may also be linked to the change in dominant strains as the season progresses, potentially implicating nutrient limitation as a driver of strain succession. Associations between cyanobacteria colonies and attached vorticellids (Figure 9) may relate to mobility and preferred water column depth, as the combination of cyanobacterial gas vacuoles and the protozoan’s enhanced buoyancy and water movement could enhance the movement of the cyanobacterium colony through the water (Canter et al., 1992).

**Figure 9.**
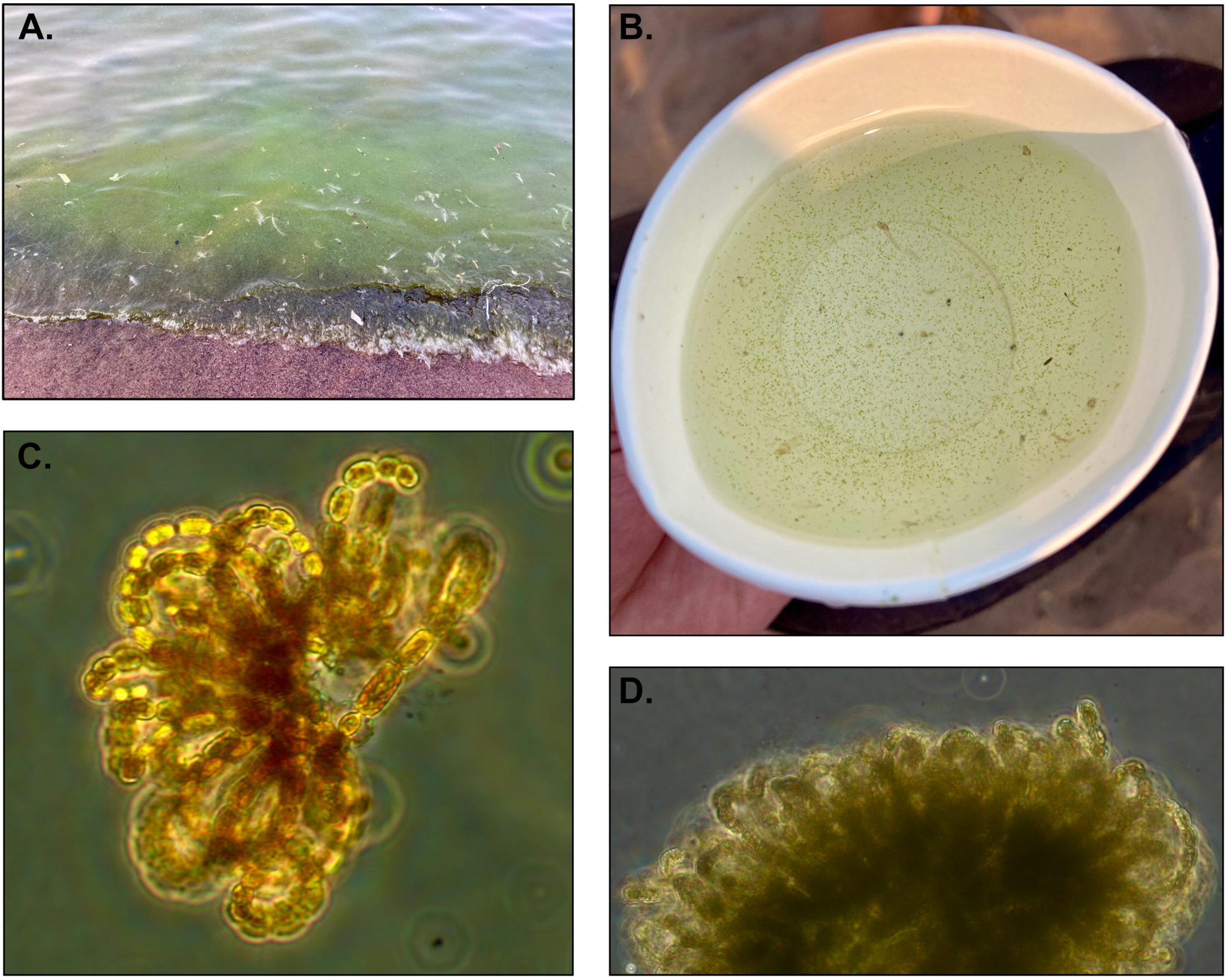
The presence of several *Vorticella sp.* attached to a colony of *D. lemmermannii* from the 2022 net tow samples.

Through our comprehensive 16S rRNA gene survey, we sought to test if the abundance of *D. lemmermannii* was correlated with environmental conditions and nutrient concentrations taken during sampling. We generally detected no significant positive or negative correlations with water column conditions (Table S2). Several variables, like water column depth, distance from the river mouth, and nitrate concentrations, were weakly positively correlated with 16S ASV abundance, suggesting there may be a slight preference for offshore stations. However, the variability observed between stations and the near-shore and offshore environment suggests other factors are at play. For example, net tows in 2022 generally showed higher colony counts closer to shore than offshore sites. One possibility is that the growth of *D. lemmermannii* is not tied to the water column conditions measured at the time of sampling. As a water package moves, so does the concentration of nutrients within that water package, similar to the transport of buoyant fluids and microorganisms from hydrothermal vents (Reed et al., 2015; Sheik et al., 2014). As the organisms are co-transported, the nutrient concentrations can decrease due to microbial metabolism and mixing. Further complicating our correlations is that *D. lemmermannii* likely can store nutrients like carbon, nitrogen, phosphorus, and iron (Sheik et al., 2022). Thus, changes in nutrient concentration may be negated by the ability of *D. lemmermannii* to supplement external nutrients and maintain growth. A second possibility is that wind-driven surface currents influence the dispersion and abundance of organisms and potentially the depths at which they occur during the day. We see evidence of this at several stations where ASVs were present in the nearshore and offshore stations but not in between these stations (Figure 2). Because we took samples at a defined epilimnion depth (2 m), we may have missed populations if they were deeper. Anecdotally, blooms are often observed after a period of calm conditions, suggesting that floating and then drifting is an understudied but important process.

The presence of multiple, differentially abundant genome strains within Lake Superior also highlights that strains may specialize in specific water column conditions. This would suggest that past blooms across Lake Superior may be driven by different nutrient or environmental conditions depending on the timing and potentially the location of the blooms. Nonetheless, time is a key factor for determining where and when these blooms initiate and ultimately manifest. Using gRodon (Weissman et al., 2021), which estimates the growth rate of microorganisms based on their genome, we predict the doubling time of *D. lemmermannii* under optimal, nutrient-replete conditions is around 19-20 hours. There are caveats to using this methodology (Weissman et al., 2021), and it is well-known that growth estimates are difficult for slow-growing organisms (Gonzalez and Aranda, 2023). That said, to form a colony of *D. lemmermannii* with 500 cells from an akinete, would take ∼7 days under optimal conditions. Microorganisms rarely encounter optimal conditions, particularly in Lake Superior, where low to moderate temperatures and limitations of phosphorus, iron, or manganese will greatly affect the growth rates of cyanobacteria. Expected doubling times in situ would therefore be longer than under ideal conditions. Furthermore, these nutrients’ limitations have likely contributed to the accumulation of nitrate in Lake Superior (Sterner et al., 2020). Thus, under suboptimal conditions, *D. lemmermannii* would need days to weeks to develop enough biomass to create a bloom. Pitcher et al. (2010) describe how various coastline features such as islands and bays impact hydrological factors like retention time that, in turn, influence the development of harmful algal blooms. It is possible that water currents interact with the Apostle Islands and the series of bays starting at Bark Point to increase retention times and concentrate *D. lemmermannii*, alter local nutrient availability, influence water temperatures, or some combination of these factors, and thereby provide the conditions necessary to accumulate sufficient biomass for bloom formation. Our data are consistent with this hypothesis, as we observe high relative abundances of *D. lemmermannii* at sampling stations in the area surrounding Siskiwit Bay on July 20th in both our 16S and metagenomic data and again in late September in the 16S dataset (Figure 2, Figure 5). Interestingly, a similar pattern is apparent in our net tow data the following year, with elevated abundances but relatively low colony density at the eastern, near-shore sites throughout the season (Figure 6). Already notable as a bloom hotspot over the past decade, these high relative abundances of *D. lemmermannii* under non-bloom conditions in the area surrounding Siskiwit Bay certainly suggests that something about this region is especially amenable to *D. lemmermannii* growth and blooms. Better models of water movement for this part of Lake Superior are necessary to interrogate this hypothesis, but little relevant data exists. Future studies to close this gap would be useful to pair with sequence-based monitoring of *D. lemmermannii* population dynamics. Additionally, recurring sequences in the sediment (Sheik et al., 2022) reinforce the conclusion that *D. lemmermannii* is well-established in the microbial community and not an ephemeral species dependent on periodic influxes from inland source populations.

## Conclusions

This study finds that *D. lemmermannii* is widely distributed under non-bloom conditions along the Wisconsin shore of Lake Superior between Duluth and the Apostle Islands. We observe spikes in the relative abundance of this bloom-forming cyanobacterium in the area of Siskiwit Bay in mid-summer and early fall, highlighting that location as one with a high risk of bloom formation. Managers may use this insight to focus monitoring efforts when this risk is highest. Genome-resolved analysis showed a striking pattern of *D. lemmermannii* strain succession throughout the summer, suggesting that different strains are adapted to changing environmental conditions. It remains unclear what environmental variables may be driving strain turnover or the spatiotemporal patterns of abundance that we observe. Future studies should expand on this work in several ways. First, a higher frequency of sampling would improve the temporal resolution, and coupling this to an expanded sampling area, such as the north shoreline and watershed would shed light on the geographic provenance of different strains and may capture *in situ* growth of *D. lemmermannii.* Additionally, the addition of high-resolution depth profiles, instead of just the surface, may reveal *D. lemmermannii* in places we do not expect, revealing other variables that structure its population and distribution. Deeper metagenomic sequencing would also permit higher-resolution population genomic comparisons and facilitate comparative genomic analyses to identify ecological adaptations pertinent to strain succession and bloom formation (e.g., temperature adaptation). Finally, coupling growth and water transport models could facilitate new views of microbial population dynamics in these large lakes that may not be feasible using traditional sampling methods.

## Acknowledgments

We thank the BRICO Fund and National Park Service for providing funding to make the sampling effort and amplicon sequencing component of this work possible. The National Park Service also provided funding for the net tow studies. We also thank the Cooperative Institute for Great Lakes Research (CIGLR) for providing rapid funds to perform the metagenomic component of this study. The authors have no conflicts of interest to report.

